# Determination of primary microRNA processing in clinical samples by targeted pri-miR-sequencing

**DOI:** 10.1101/2020.04.30.070912

**Authors:** Thomas Conrad, Evgenia Ntini, Benjamin Lang, Luca Cozzuto, Jesper B Andersen, Jens U Marquardt, Julia Ponomarenko, Gian Gaetano Tartaglia, Ulf A V Ørom

## Abstract

MicroRNA expression is important for gene regulation and deregulated microRNA expression is often observed in disease such as cancer. The processing of primary microRNA transcripts is an important regulatory step in microRNA biogenesis. Due to low expression level and association with chromatin primary microRNAs are challenging to study in clinical samples where input material is limited.

Here, we present a high-sensitivity targeted method to determine processing efficiency of several hundred primary microRNAs from total RNA using as little as 500 thousand Illumina HiSeq sequencing reads. We validate the method using RNA from HeLa cells and show the applicability to clinical samples by analyzing RNA from normal liver and hepatocellular carcinoma.

We identify 24 primary microRNAs with significant changes in processing efficiency from normal liver to hepatocellular carcinoma, among those the highly expressed miRNA-122 and miRNA-21, demonstrating that differential processing of primary microRNAs is occurring and could be involved in disease. With our method presented here we provide means to study pri-miRNA processing in disease from clinical samples.

## Introduction

MicroRNAs (miRNA) are small RNAs that regulate gene expression at the post-transcriptional level [1]. miRNAs are transcribed as primary miRNA (pri-miRNA) that can be several kilobases long. These transcripts are processed in the nucleus to 60-90 nts long precursor miRNA hairpins (pre-miRNA) by the Microprocessor complex. These pre-miRNAs are subsequently exported to the cytoplasm by export factors where they are processed into 20-23 nts long mature miRNAs by Dicer and incorporated into the RISC (RNA-induced silencing complex) to exert their regulatory function [1].

We have previously showed that the endogenous Microprocessor activity toward individual pri-miRNAs can be determined using RNA sequencing [2]. We identified a Microprocessor cleavage signature and defined a metric for processing efficiency. We showed on a transcriptome-wide scale that processing efficiency is highly variable among different canonical pri-miRNAs and a major determinant of the expression levels of individual mature miRNAs. In addition, we have showed that the kinetics of pri-miRNA processing vary between individual transcripts, including within polycistronic transcripts [3].

*In vitro* assays looking at individual pri-miRNA transcripts have shown pri-miRNA processing to be an actively regulated process that is responsive to TGFß signaling [4], DNA damage [5] and cell density [6], and a recent study applied our pri-miRNA processing approach to demonstrate that activation of the Type I interferon response in cells affects general pri-miRNA processing [7]. Several Microprocessor co-factors have been identified that influence pri-miRNA processing [2] and composition of the sequence flanking the pre-miRNA hairpin have been shown to affect efficiency of Microprocessor cleavage [2, 8].

The major limitation of our approach to profile pri-miRNA processing transcriptome-wide has been a requirement for purification of chromatin-associated RNA and large sequencing depth, making analysis of clinical samples unfeasible. In the work reported here we aimed to improve the approach to enable the analysis of pri-miRNA processing in clinical samples directly from whole cell total RNA isolated from tissue. We used a targeted sequencing approach allowing for quantification of 361 pri-miRNAs with as little as 500 thousand Illumina HiSeq reads, and demonstrate the applicability to clinical samples using RNA from normal liver (NL) and hepatocellular carcinoma (HCC) samples. We also identified differentially processed pri-miRNAs between NL and HCC with a potential implication in disease.

## Results

To determine pri-miRNA processing experimentally in clinical samples we expanded our RNA-sequencing based methodology [2]. As mentioned above, the mature miRNA is generated in a sequential manner starting with the transcription of a primary transcript [1]. The pre-miRNA is cut out of the pri-miRNA by the Microprocessor complex (Figure 1a) consisting of Drosha and DGCR8, in addition to a number of co-factors [9]. When using RNA-sequencing to high depth of chromatin-associated RNA (enriching for pri-miRNA transcripts) a Microprocessor signature is observed where the pre-miRNA has been cut out (Figure 1b) [2, 3]. This signature can be quantified by determining the ratio between reads covering the pre-miRNAs and the flanking sequence on the pri-miRNAs [2] (Supplementary Figure 1).

**Figure 1.**
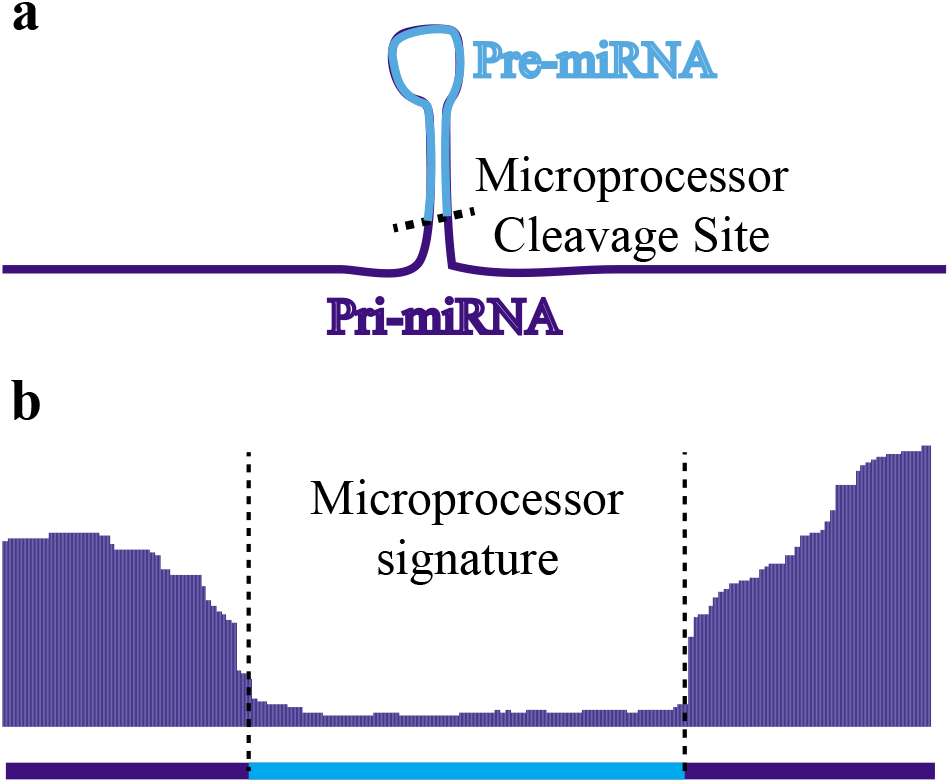
The Microprocessor signature. (a) The general structure of a pri-miRNA with the hairpin pre-miRNA and the Microprocessor cleavage site indicated. (b) Example of the Microprocessor signature around the pre-miRNA in an RNA sequencing read density plot.

The major limitation of our published methodology is the requirement for enrichment of chromatin-associated RNA and a high sequencing depth (for the development of the method we used 200 million reads per sample). These requirements made the analysis of pri-miRNA processing in total RNA from tissue and clinical samples unfeasible.

To overcome this limitation we developed a targeted approach to determine processing efficiency of a library of selected pri-miRNAs from small amounts of starting material of total RNA and low sequencing depth. We designed enrichment probes (xGEN lockdown probes, Integrated DNA Technologies) covering the sequence of the pri-miRNA flanking the pre-miRNA hairpin both upstream and downstream (Figure 2a).We targeted 32 pri-miRNAs that we had already determined the processing efficiency for using chromatin-associated RNA and high sequencing depth [2]. Each probe was designed to be complementary to the 120 nucleotides immediately upstream or downstream of the Microprocessor cleavage sites, respectively. We performed two independent enrichment experiments on RNA sequencing libraries generated from chromatin-associated RNA to test the reproducibility of the enrichment approach (Figure 2b). We see a highly reproducible processing efficiency across all 32 pri-miRNAs for the two samples (R = 0.995), demonstrating that the approach is robust and generates reproducible data. We next asked how well the enrichment approach recapitulates the processing efficiencies determined by high-depth sequencing without enrichment (Data from Conrad et al., 2014). From both independent probe enrichment replicates we see a high correlation (R = 0.92) in processing efficiency when comparing to previous data of pri-miRNA processing from purified chromatin-associated RNA (Figure 3a) [2]. Processing efficiencies and coverage for the 32 pri-miRNAs are shown in Supplementary Table 1.

**Figure 2.**
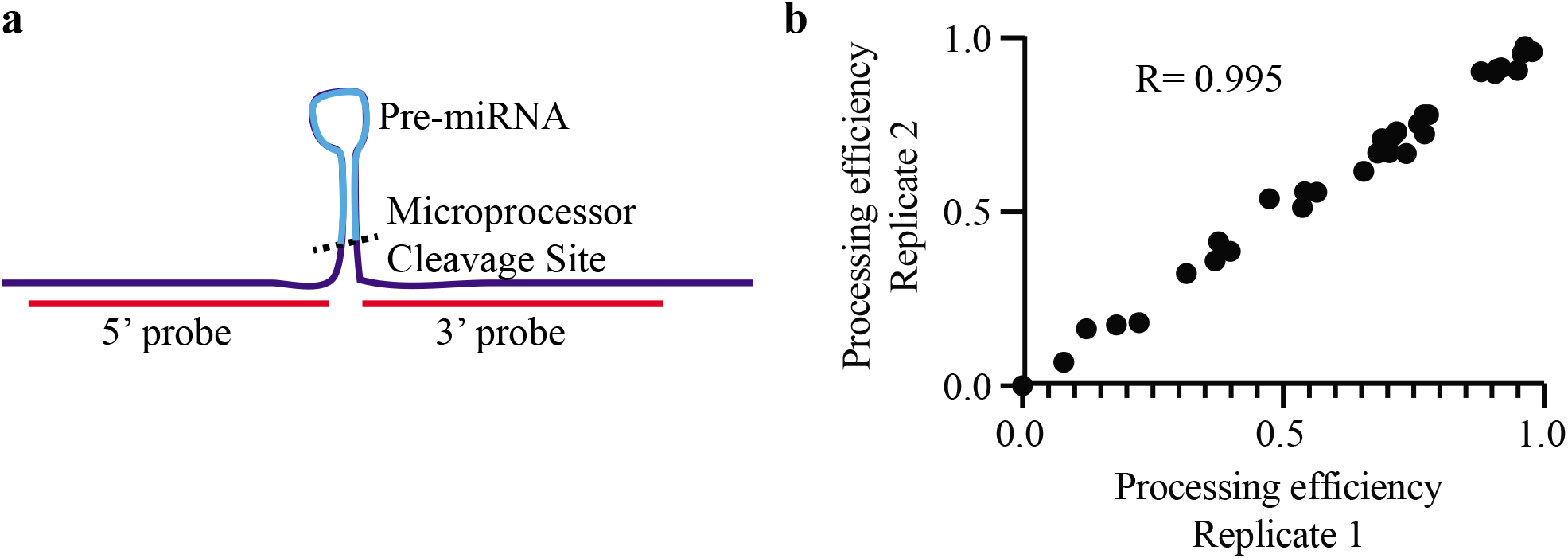
Enrichment of pri-miRNAs for determination of processing efficiency. (a) Schematic of a pri-miRNA and the localization of our enrichment probes. (b) Reproducibility of processing efficiency for 32 pri-miRNAs between two independent replicates from RNA sequencing libraries generated from chromatin-associated RNA.

**Figure 3.**
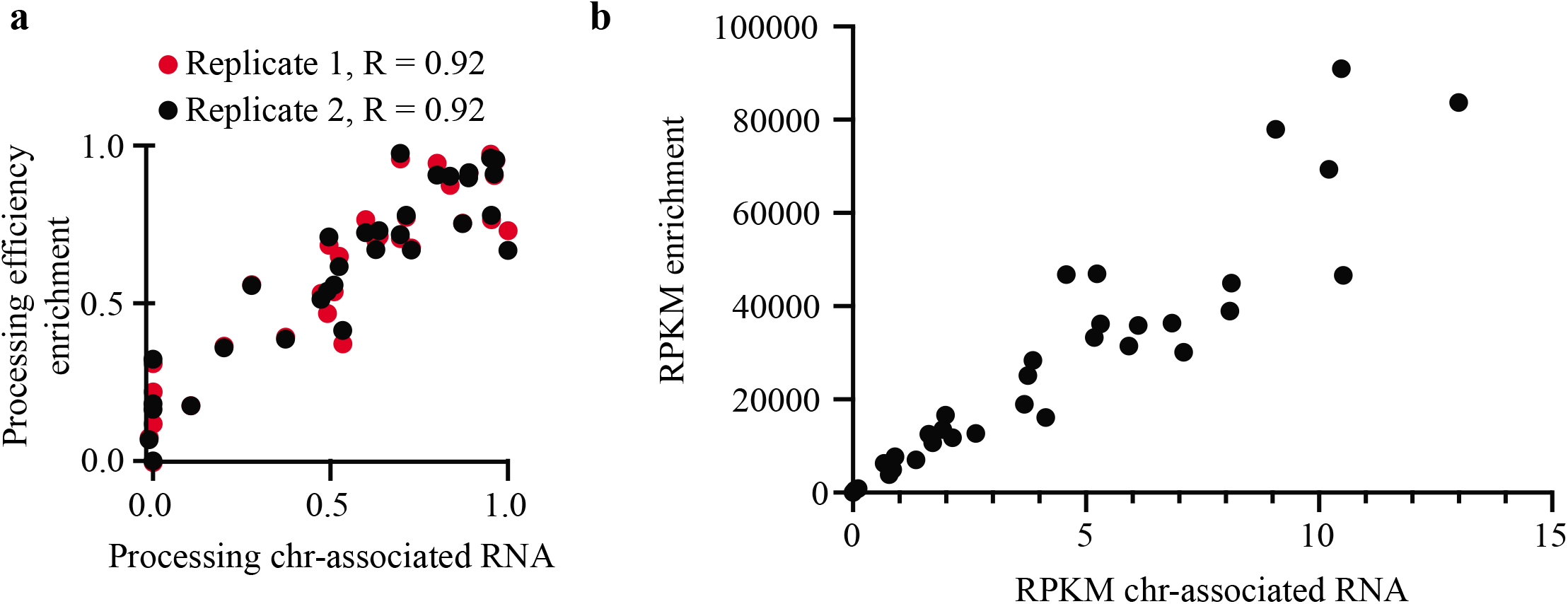
Reproducibility and sensitivity of pri-miRNA processing efficiency determination. (a) Correlation of processing efficiency determined from high-depth sequencing of chromatin-associated RNA [2] and low-depth sequencing of targeted sequencing of total RNA for two technical replicates. (b) Sensitivity of each approach determined by RPKM at pri-miRNAs. Slope shows a 6,670-fold increase in sensitivity and a corresponding decreased need for sequencing depth. RPKM for targeted sequencing is calculated as the average of two independent enrichment replicates.

To assess the degree of enrichment achieved with the targeted approach we calculated RPKM (Reads Per Kilobase per Million reads) for each of the 32 pri-miRNAs. As shown in Figure 3b the enrichment is linear across the expression range of assayed pri-miRNAs (R = 0.998) with a slope of 6,670-fold enrichment when using targeted sequencing of pri-miRNA transcripts. This means that, in principle, 30,000 reads should be sufficient to determine processing efficiency for selected pri-miRNAs when enriched from the chromatin fraction. Assuming that chromatin-associated RNA constitutes 5 per cent of the total cellular (non-ribosomal) RNA 600,000 reads is necessary to achieve robust determination of pri-miRNA processing from total RNA after targeted enrichment.

To address differential pri-miRNA processing in clinical samples we used RNA from 40 HCC tumors and 9 NL samples. We selected 361 miRNAs with a described involvement in cancer [10–12] and designed 120-nts xGEN enrichment probes as illustrated in Figure 2a. For each RNA sample we prepared a library for Illumina HiSeq 2500 sequencing and enriched with the probe library as described in Materials and Methods. We aimed for 1 million reads per library when multiplexing and the final output per library was between 500 thousand and 3 million reads. With this sequencing depth we could detect the processing efficiency for 209 pri-miRNAs in at least two samples, and show differential processing between HCC and NL for 24 of the pri-miRNAs included in the analysis (P < 0.05, Wilcoxon rank test) (Figure 4a and Supplementary Table 2). Of particular interest, we see the most statistically significant changes in the processing of the liver-specific miR122 and the oncomiR miR21 (Figure 4b), where processing becomes more efficient in HCC. In NL around 20 per cent of the pri-miRNA remains unprocessed whereas in HCC there is an almost complete processing (a few per cent of the pri-miRNAs remain unprocessed).

**Figure 4.**
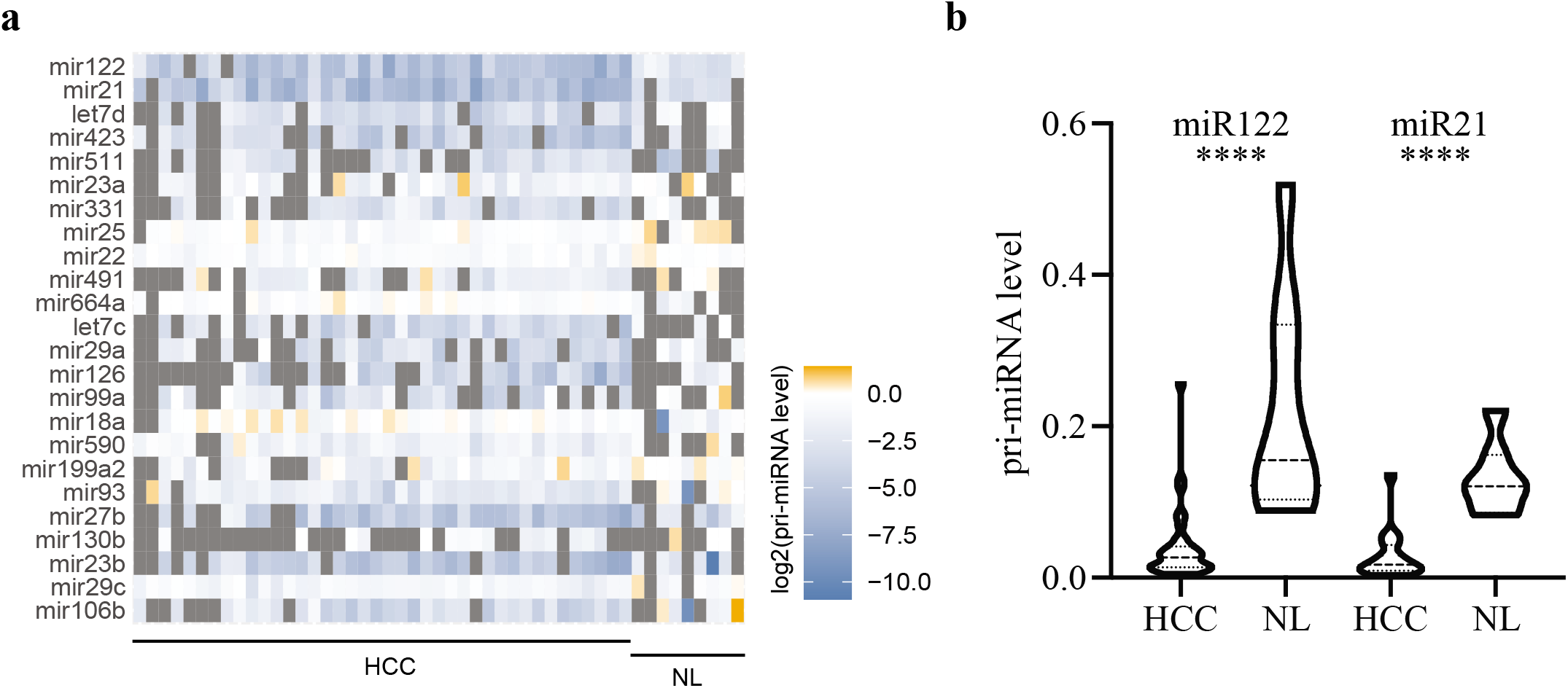
Differential pri-miRNA processing between NL and HCC. (a) Heat-map of pri-miRNA levels of the 24 differentially processed pri-miRNAs between HCC and NL. (b) Increased processing of pri-miRNAs in HCC compared to NL for miR-122 and miR-21. Shown is the pri-miRNA level as the proportion of the pri-miRNA that remains unprocessed as determined by RNA sequencing. **** p < 0.0001, Wilcoxon rank test.

## Discussion

We have previously shown that the processing efficiency of individual pri-miRNAs is similar between three different cell lines (HeLa, HEK293 and A549), suggesting that diversity in processing is largely dictated by the diverse substrate sequences. On the other side, several studies have shown that co-factors and signaling pathways can regulate processing of individual pri-miRNAs or groups of pri-miRNAs [4–8].

Here, we have developed a targeted version of our approach to measure endogenous pri-miRNA processing for hundreds of different pri-miRNA at a time. Our targeted approach offers high sensitivity to the selected pri-miRNAs and allows assessing numerous pri-miRNAs at a time in a large collection of tissue and clinical samples. The method is particularly feasible to complement total RNA sequencing studies as existing RNA sequencing libraries can directly be used for targeted sequencing of pri-miRNA transcripts at very low cost once the enrichment probe library has been designed.

We show that the targeted approach works well for clinical samples by demonstrating reproducible differential processing of a number of pri-miRNAs between NL and HCC. Here, we show that differential pri-miRNA processing occurs between NL and HCC suggesting that the primary processing step in miRNA biogenesis can impact gene expression in disease. Our targeted approach offers the possibility for further studies of pri-miRNA processing from clinical samples that has so far not been feasible, allowing identification of the molecular consequences of deregulated pri-miRNA processing in disease.

## Methods

### Design of capture probes

Capture probes were designed as xGEN Lockdown probes from IDT with the minimal recommended length of 120 nts. For each pri-miRNA targeted by the library we made two probes, one upstream and one downstream of the Microprocessor processing site.

### Enrichment of RNA sequencing library

Targeted RNA sequencing libraries were enriched according to the xGEN guidelines with minor modifications. In brief, 12 barcoded RNA sequencing libraries were pooled and 500 ng used for each target enrichment. Blocking oligos were added to the pooled libraries, samples were dried in a SpeedVac and resuspended in Hybridization buffer including the custom enrichment probes and hybridized at 65 degrees for 4 hours after a 30 second incubation at 95 degrees. Streptavidin beads were washed twice in Bead Wash Buffer before the capture reaction of custom enrichment probes and targeted sequencing library. Samples were incubated 45 minutes at 65 degrees followed by normal wash and two times high-stringency heated washes followed by an additional wash at room temperature and elution of captured library with nuclease-free water. The captured libraries were amplified with PCR using KAPA HiFi Hotstart polymerase for 14 cycles. The post-capture PCR fragment were purified with Agencourt AMPure XP beads before sequencing of the targeted libraries.

### Data analysis and calculation of processing efficiency

Raw reads were inspected for quality using FastQC (v0.11.5). Reads were then trimmed with skewer (version 0.2.2) for removing the adapters and low quality reads. Processed reads were aligned to the reference genome (human, GENCODE release 27, GRCh38.p10) using STAR aligner (version 2.5.3a). bedtools “genomecov” was run on the aligned reads with the “-split” option for calculating the coverage. The coverage was multiplied for a scaling factor (using the parameter “-scale”) that is obtained by dividing 1 billion / (number of mapped reads * read size). The BED graph was converted to bigWig using bedGraphToBigWig. Custom scripts were used to calculate read count averages and the processing efficiency was calculated by counting the reads covering the pre-miRNA region divided by the reads covering the two 100 nts flanking regions of the pri-miRNA (starting 20 nts from the stem to avoid noise in read coverage), and normalized to the length of each pri-miRNA in nts (Supplementary Figure 1). Pre-miRNAs were annotated from the sequence of the mature miRNAs so that each pre-miRNA starts with the first base of the 5P miRNA and ends with the last base of the 3P miRNA.

### Data availability

All sequencing data have been deposited to GEO under accession number GSE148756 and GSE149631.

## Author contributions

Conceived experiments: TC, UAVØ; Performed experiments: TC, UAVØ; Analyzed data: EN, BL, LC; Contributed clinical samples: JBA, JM; Supervised computational work: EN, JP, GGT; Supervised experimental work: UAVØ; Interpreted data: TC, EN, BL, LC, JP, GGT, UAVØ; Secured funding: UAVØ; Wrote the manuscript: TC, BL, UAVØ; Commented on the manuscript and approved the final version: All authors.

## Acknowledgments

This work was funded by the Sofja Kovalevskaja Award of the Alexander von Humboldt Foundation and the Hallas Møller Award of the Novo Nordisk Foundation (UAVØ).

