## Supplementary data for "Determination of primary microRNA processing in clinical samples by targeted pri-miR-sequencing"

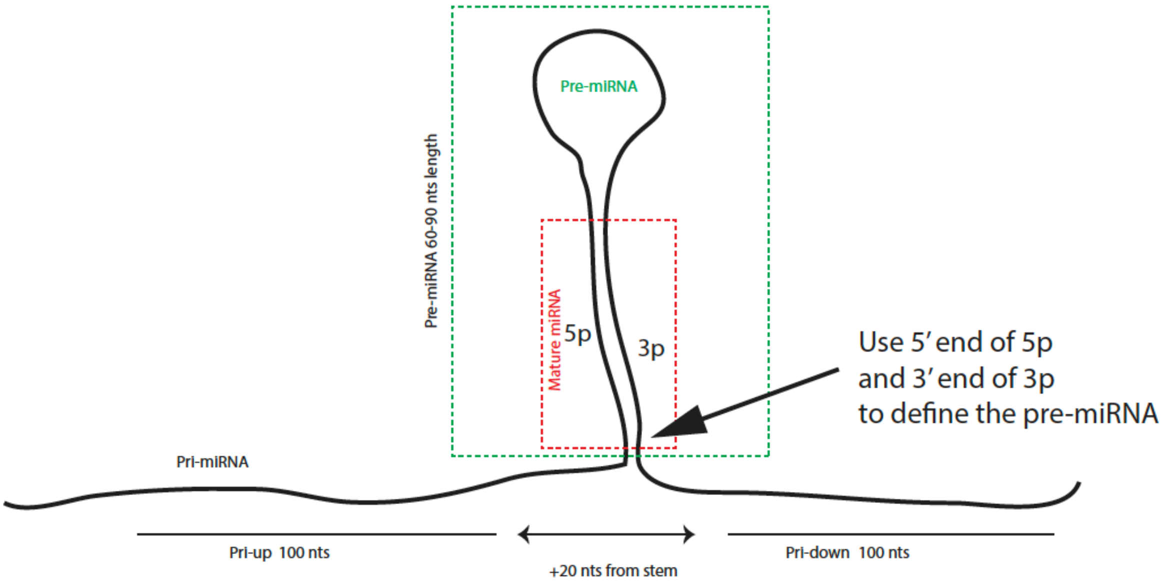


$$\frac{\text{Reads (pre-miRNA)}}{\text{Reads (Pri-up + Pri-down)}} * \frac{200}{\text{Length of pre-miRNA}}$$

**Supplementary Figure 1**

Overview of the pri-miRNA organization and how the processing efficiency is calculated.

**Supplementary Table 1**

Processing efficiency and RPKM for targeted enrichment of 32 pri-miRNAs determined in HeLa cells. Shown are also PE and RPKM for total chromatin-associated RNA from Conrad et al., 2014.

**Supplementary Table 2**

Processing efficiency of targeted enrichment of 361 pri-miRNAs for 40 HCC samples and 9 Normal Liver samples. 361 pri-miRNAs are targeted with the custom probe library, of these, 209 are detected in more than one sample.
