## Supplementary Table 1 for "Determination of primary microRNA processing in clinical samples by targeted pri-miR-sequencing"

| id | HeLa rep A PE | HeLa rep B PE | Chr PE | RPKM rep A | RPKM rep B | RPKM Chr |
|---|---|---|---|---|---|---|
| hsa-let-7a-1 | 0.918 | 0.914 | 0.891 | 69440 | 69392 | 10 |
| hsa-let-7a-3 | 0.880 | 0.903 | 0.838 | 30987 | 31968 | 6 |
| hsa-let-7b | 0.542 | 0.559 | 0.511 | 35302 | 37157 | 5 |
| hsa-let-7d | 0.537 | 0.514 | 0.474 | 45800 | 44049 | 8 |
| hsa-let-7f-1 | 0.690 | 0.711 | 0.496 | 35952 | 36801 | 7 |
| hsa-let-7i | 0.315 | 0.323 | 0.000 | 38775 | 39170 | 8 |
| hsa-mir-106b | 0.760 | 0.753 | 0.872 | 46068 | 47108 | 11 |
| hsa-mir-10b | 0.963 | 0.976 | 0.697 | 822 | 981 | 0 |
| hsa-mir-125a | 0.950 | 0.907 | 0.801 | 6413 | 6176 | 1 |
| hsa-mir-181a-1 | 0.958 | 0.956 | 0.966 | 19334 | 18507 | 4 |
| hsa-mir-181b-1 | 0.080 | 0.068 | -0.011 | 13024 | 12377 | 3 |
| hsa-mir-197 | 0.181 | 0.176 | 0.108 | 12726 | 12403 | 2 |
| hsa-mir-21 | 0.779 | 0.779 | 0.714 | 860460 | 863574 | 143 |
| hsa-mir-218-1 | 0.736 | 0.669 | 1.000 | 493 | 640 | 0 |
| hsa-mir-221 | 0.718 | 0.731 | 0.637 | 35602 | 36163 | 6 |
| hsa-mir-222 | 0.910 | 0.911 | 0.962 | 27929 | 28834 | 4 |
| hsa-mir-23a | 0.398 | 0.387 | 0.374 | 24491 | 25706 | 4 |
| hsa-mir-23b | 0.710 | 0.718 | 0.697 | 47590 | 46287 | 5 |
| hsa-mir-24-1 | 0.565 | 0.557 | 0.279 | 32836 | 33761 | 5 |
| hsa-mir-24-2 | 0.224 | 0.182 | 0.000 | 13123 | 13955 | 2 |
| hsa-mir-25 | 0.000 | 0.000 | 0.000 | 83006 | 84404 | 13 |
| hsa-mir-27a | 0.473 | 0.538 | 0.492 | 16042 | 17200 | 2 |
| hsa-mir-27b | 0.370 | 0.360 | 0.200 | 78551 | 77300 | 9 |
| hsa-mir-28 | 0.377 | 0.414 | 0.535 | 6892 | 7154 | 1 |
| hsa-mir-335 | 0.978 | 0.961 | 0.952 | 3964 | 3779 | 1 |
| hsa-mir-374a | 0.771 | 0.725 | 0.599 | 11815 | 11847 | 2 |
| hsa-mir-378a | 0.681 | 0.669 | 0.729 | 7731 | 7730 | 1 |
| hsa-mir-423 | 0.906 | 0.898 | 0.889 | 91876 | 90122 | 10 |
| hsa-mir-545 | 0.123 | 0.165 | 0.000 | 10696 | 10691 | 2 |
| hsa-mir-652 | 0.770 | 0.780 | 0.954 | 5017 | 4849 | 1 |
| hsa-mir-93 | 0.704 | 0.671 | 0.629 | 46760 | 46863 | 5 |
| hsa-mir-99a | 0.654 | 0.616 | 0.525 | 15642 | 16532 | 4 |

Chr data are from Conrad et al., 2014
PE - Processing Efficiency
RPKM - Reads Per Kilobase per Million reads
